# Data gaps of international databases on HPAI H5 in wildlife in the Americas: implications for surveillance, research, and conservation

**DOI:** 10.64898/2026.05.30.728949

**Authors:** Ralph E. T. Vanstreels, Marcela M. Uhart

## Abstract

Global efforts to prevent and mitigate the impacts of high pathogenicity avian influenza (HPAI) H5 on domestic animals, humans, and wildlife rely on timely and transparent information that is both accurate and interpretable across countries and sectors. International epidemiological and genomic databases, such as the World Animal Health Information System (WAHIS), the Global Animal Disease Information System (EMPRES-i+), the Global Initiative on Sharing All Influenza Data (GISAID), and the National Center for Technological Bioinformation Virus Portal (NCBI) provide essential information for surveillance, research, and decision-making. To evaluate how well these resources capture recent wildlife impacts, we consolidated information from these databases and complementary public sources including government reports, scientific literature, and news articles, on wildlife mortality associated with HPAI H5 in the Americas from November 2021 to July 2024. The consolidated dataset comprised 615,883 wild birds (287 spp.) and 63,409 wild mammals (39 spp.). In comparison, WAHIS represented 16,902 wild birds (261 spp.) and 6,323 wild mammals (31 spp.) while EMPRES-i+ captured a substantially smaller portion of affected host diversity for both wild birds (105 spp.) and wild mammals (27 spp.). Genomic databases (GISAID and NCBI) represented 7,027 whole genome equivalents of H5 viruses from wild birds (175 spp.) and 371 from wild mammals (26 spp.). These discrepancies indicate that international databases, while essential, provide an incomplete picture of HPAI impacts on wildlife, with significant geographic and taxonomic asymmetries attributable to differences in surveillance capacity, reporting practices, sequencing effort, and data-sharing pathways. Studies and management strategies relying on these resources without complementary validation may therefore mistake data gaps for real-world epidemiological patterns. Strengthening data reporting standards, improving validation procedures, and integrating international databases with national reports, scientific publications, and other sources will enhance the reliability of epidemiological analyses and support more effective One Health surveillance, risk assessment, and conservation action.

**Author summary:** High pathogenicity avian influenza (HPAI) H5 viruses, often called bird flu viruses, can cause severe disease in birds and mammals, including humans. Because of their relevance for human health, livestock production, and wildlife conservation, international databases were established to share information on when and where these viruses are detected, which species are affected, and what virus strains are found. These databases are essential tools for governments, scientists, and conservation practitioners working to track outbreaks, understand how these viruses spread and evolve, and guide surveillance and response.

In this study, we compiled and compared information on recent HPAI H5 events in wildlife in the Americas available in international databases with information from other public sources, including reports from governments, scientific literature, and news articles. We found important discrepancies in how countries and species affected were represented across sources. As a result, international databases might not fully capture the actual distribution or conservation impact of HPAI H5 on wildlife. Our findings also show why decision-makers and scientists should interpret database-derived patterns carefully. We provide recommendations to improve international databases to address these gaps and better inform One Health risk assessment and wildlife conservation actions.

## Introduction

The global spread of high pathogenicity avian influenza (HPAI) H5 viruses of the Eurasian Goose/Guangdong lineage since 2021 has raised significant concern due to its impacts for the poultry and dairy cattle industry, human health and livelihoods, and biodiversity conservation [1,2]. Early in the panzootic, European seabird populations were heavily impacted with some species experiencing population decreases as severe as 25–70% following the 2021–2022 wave of HPAI H5 [3–7]. The subsequent arrival of these viruses in North America, beginning in November 2021, marked a major expansion in wildlife in the Americas, with rapid spread among waterfowl, seabirds, and raptors [8–11] and, to a lesser extent, terrestrial and marine mammals [12,13]. By late 2022, HPAI H5 had reached Central and South America, where large-scale outbreaks in seabirds and marine mammals represented a major concern for wildlife conservation in the region [14–19].

Global efforts to prevent, detect, and mitigate the impacts of HPAI H5 in domestic animals, humans and wildlife rely on timely and transparent information that is both accurate and interpretable across countries and sectors. The World Animal Health Information System (WAHIS) is a database built to address this need, providing information on global animal health status as reported to the World Organisation for Animal Health (WOAH) by its member states [20]. Because HPAI is a WOAH-notifiable disease, WAHIS is widely recognized by animal health researchers and stakeholders as a key source of information on the global distribution of HPAI outbreaks. This is illustrated by recent articles relying on WAHIS data to evaluate the conservation impacts, spatiotemporal dissemination patterns, correlation to environmental variables, transmission patterns, and species susceptibility, among other aspects of HPAI H5 ecology [21–28]. Likewise, the Global Animal Disease Information System (EMPRES-i+) is a database maintained by the Food and Agriculture Organization of the United Nations (FAO) that collects information from various sources –including WAHIS, national authorities, and FAO officers– on animal disease outbreaks, including HPAI [29,30]. EMPRES-i+ aims to provide assistance to veterinary services for their disease control activities, and its data has also been used in HPAI epidemiological and phylodynamic research [24,30–32]. Comparably, the Global Initiative on Sharing All Influenza Data (GISAID) and the National Center for Technological Bioinformation Virus Portal (NCBI) are large databases for virus genomic data. These two genomic databases have been extensively used for research and surveillance, including the detection of new variants and reassortment events, monitoring of evolutionary patterns and genotype competition, inferring on the phenotypic, epidemiological, and risk consequences of mutations, identifying vaccine candidate strains, and predicting pandemic potential, among others [33–39]. Together, these resources provide critical context for understanding HPAI H5 ecology, evolution, and impacts at regional and global scales.

Information derived from these databases may be influenced by surveillance capacity, reporting practices, sequencing effort, and public data-sharing pathways across countries and host taxa [33,40]. This has important implications for One Health surveillance, risk assessment and conservation planning, particularly when wildlife mortality is extensive but unevenly documented. To assess these potential issues, we compiled and compared data from WAHIS, EMPRES-i+, GISAID, and NCBS with complementary public sources on wildlife mortality associated with HPAI H5 in the Americas from November 2021 to July 2024. The gaps and asymmetries we uncovered in the representation of wildlife mortality due to HPAI H5 across these data sources highlight opportunities to improve how these data are used in surveillance, research, risk assessment, and conservation planning.

## Results

### Overall publicly-reported wildlife mortality associated with HPAI H5

Our consolidated dataset based on public information (S1 File) amounts to 615,883 wild birds (287 species, 54 families) and 63,409 wild mammals (39 species, 15 families) found dead or estimated to have died in association with HPAI H5 in the Americas during the study period. Reported mortality was predominantly concentrated in South America, with 519,865 wild birds and 62,600 wild mammals, followed by Northern America with 95,165 wild birds and 809 wild mammals, and Central America and the Caribbean with 853 wild birds. Wild bird mortality was largely concentrated in Peru (421,472 individuals; 68.4% of wild bird mortality), followed by the Falkland/Malvinas Islands (53,412 individuals; 8.7%) and the United States (50,352 individuals; 8.2%), whereas wild mammal mortality was concentrated in Argentina (29,506 individuals; 46.5% of wild mammal mortality), Chile (20,288 individuals; 32.0%), and Peru (9,025 individuals; 14.2%) (Fig 1 and Table 1). In relation to host taxonomy, publicly-reported mortality involved predominantly wild birds of families Phalacrocoracidae (207,990 individuals; 33.8% of wild bird mortality), Sulidae (197,827 individuals; 32.1%), and Pelecanidae (55,947 individuals; 9.1%) and wild mammals of families Otariidae (35,424 individuals; 55.9% of wild mammal mortality) and Phocidae (27,375 individuals; 43.2%) (Fig 2 and Table 2).

**Table 1.** Summary of the distribution of HPAI H5 in wildlife in different datasets among countries/territories in the Americas. Genomic data availability was quantified in whole genome equivalents (number of gene segment sequences divided by eight). The denominator of the representation ratio was calculated dividing the publicly-reported mortality (number of individuals) by the WAHIS-reported affected wildlife (number of individuals) or by the available H5 virus genomic data (whole genome equivalents). Study period: November 2021 – July 2024 (inclusive).

| Country/territory | Number of individuals |  | Whole genome equivalents |  |  | Denominator of the representation ratio |  |  |  |
| --- | --- | --- | --- | --- | --- | --- | --- | --- | --- |
|  | This study | WAHIS | GISAID | NCBI | GISAID+NCBI | WAHIS | GISAID | NCBI | GISAID+NCBI |
| **Wild birds** |  |  |  |  |  |  |  |  |  |
| Argentina (ARG) | 3,005 | 476 | 6 | 4.3 | 6 | 6.3 | 500.8 | 698.8 | 500.8 |
| Bolivia (BOL) | 4 | 4 | 0 | 0 | 0 | 1 | ∞ | ∞ | ∞ |
| Brazil (BRA) | 1,172 | 1,172 | 28 | 24 | 28 | 1 | 41.9 | 48.8 | 41.9 |
| Canada (CAN) | 44,809 | 2,289 | 2,242.3 | 0 | 2,242.3 | 19.6 | 20 | ∞ | 20 |
| Chile (CHL) | 32,548 | 1,142 | 84.4 | 107.8 | 116.8 | 28.5 | 385.6 | 301.9 | 278.7 |
| Colombia (COL) | 734 | 388 | 1 | 0 | 1 | 1.9 | 734 | ∞ | 734 |
| Costa Rica (CRI) | 12 | 12 | 4.5 | 0 | 4.5 | 1 | 2.7 | ∞ | 2.7 |
| Cuba (CUB) | 39 | 34 | 0 | 0 | 0 | 1.1 | ∞ | ∞ | ∞ |
| Ecuador (ECU) | 7,181 | 7,019 | 3.9 | 0 | 3.9 | 1 | 1,841.3 | ∞ | 1,841.3 |
| Falkland/Malvinas (FLK) | 53,412 | 443 | 2 | 0 | 2 | 120.6 | 26,706 | ∞ | 26,706 |
| Greenland (GRL) | 4 | 4 | 2 | 0 | 2 | 1 | 2 | ∞ | 2 |
| Guatemala (GTM) | 11 | 11 | 2 | 0 | 2 | 1 | 5.5 | ∞ | 5.5 |
| Honduras (HND) | 325 | 325 | 1 | 0 | 1 | 1 | 325 | ∞ | 325 |
| Mexico (MEX) | 451 | 447 | 0 | 1 | 1 | 1 | ∞ | 451 | 451 |
| Panama (PAN) | 15 | 15 | 1 | 0 | 1 | 1 | 15 | ∞ | 15 |
| Peru (PER) | 421,472 | 2,704 | 29.3 | 21.6 | 32.6 | 155.9 | 14,384.7 | 19,512.6 | 12,928.6 |
| United States (USA) | 50,352 | 221 | 4,288.8 | 3,606.1 | 4,561.6 | 227.8 | 11.7 | 14 | 11 |
| Uruguay (URY) | 164 | 24 | 5.4 | 5.8 | 5.8 | 6.8 | 30.4 | 28.3 | 28.3 |
| Venezuela (VEN) | 173 | 172 | 15.5 | 2 | 15.5 | 1 | 11.2 | 86.5 | 11.2 |
| **Total** | **615,883** | **16,902** | **6,716.9** | **3,772.5** | **7,026.9** | **36.4** | **91.7** | **163.3** | **87.6** |
| **Wild mammals** |  |  |  |  |  |  |  |  |  |
| Argentina (ARG) | 29,506 | 4,758 | 14.9 | 13.5 | 15.9 | 6.2 | 1,980.3 | 2,185.6 | 1,855.7 |
| Brazil (BRA) | 1,017 | 1,017 | 4 | 4 | 4 | 1 | 254.3 | 254.3 | 254.3 |
| Canada (CAN) | 341 | 176 | 167.1 | 0 | 167.1 | 1.9 | 2 | ∞ | 2 |
| Chile (CHL) | 20,288 | 41 | 13.5 | 13.4 | 15.3 | 494.8 | 1,502.8 | 1,514 | 1326 |
| Peru (PER) | 9,025 | 3 | 7.8 | 5.8 | 7.8 | 3,008.3 | 1,157.1 | 1,556 | 1,157.1 |
| United States (USA) | 468 | 272 | 144.1 | 57 | 146.3 | 1.7 | 3.2 | 8.2 | 3.2 |
| Uruguay (URY) | 2,764 | 56 | 14.4 | 15 | 15 | 49.4 | 191.9 | 184.3 | 184.3 |
| **Total** | 63,409 | 6,323 | 365.8 | 108.6 | 371.3 | 10 | 173.3 | 583.9 | 170.8 |

**Table 2.** Summary of the apparent distribution of HPAI H5 in wildlife in different datasets among host families in the Americas. Genomic data availability was quantified in whole genome equivalents (number of gene segment sequences divided by eight). The denominator of the representation ratio was calculated dividing the publicly-reported mortality (number of individuals) by the WAHIS-reported affected wildlife (number of individuals) or by the available H5 virus genomic data (whole genome equivalents). Study period: November 2021 – July 2024 (inclusive).

| Host family | Number of individuals |  | Whole genome equivalents |  |  | Denominator of the representation ratio |  |  |  |
| --- | --- | --- | --- | --- | --- | --- | --- | --- | --- |
|  | This study | WAHIS | GISAID | NCBI | GISAID+NCBI | WAHIS | GISAID | NCBI | GISAID+NCBI |
| **Wild birds** |  |  |  |  |  |  |  |  |  |
| Accipitridae (Acc) | 1,500 | 323 | 937.5 | 570 | 952.4 | 4.6 | 1.6 | 2.6 | 1.6 |
| Alcidae (Alc) | 9,084 | 65 | 58.9 | 4 | 59 | 139.8 | 154.2 | 2271 | 154 |
| Anatidae (Ana) | 43,626 | 1,808 | 2910.5 | 1656.9 | 3102.4 | 24.1 | 15 | 26.3 | 14.1 |
| Anhimidae (Anh) | 2 | 0 | 0 | 0 | 0 | ∞ | ∞ | ∞ | ∞ |
| Apodidae (Apo) | 2 | 0 | 2 | 2 | 2 | ∞ | 1 | 1 | 1 |
| Ardeidae (Ard) | 270 | 29 | 34.8 | 17.8 | 35.8 | 9.3 | 7.8 | 15.2 | 7.5 |
| Bombycillidae (Bom) | 1 | 0 | 0 | 0 | 0 | ∞ | ∞ | ∞ | ∞ |
| Cardinalidae (Car) | 5 | 0 | 0 | 0 | 0 | ∞ | ∞ | ∞ | ∞ |
| Casuariidae (Casu) | 6 | 2 | 0.9 | 0.9 | 0.9 | 3 | 6.7 | 6.7 | 6.7 |
| Cathartidae (Cat) | 2,280 | 147 | 665 | 572.8 | 716.6 | 15.5 | 3.4 | 4 | 3.2 |
| Charadriidae (Cha) | 57 | 6 | 3 | 3 | 3 | 9.5 | 19 | 19 | 19 |
| Ciconiidae (Cic) | 2 | 1 | 2 | 2 | 2 | 2 | 1 | 1 | 1 |
| Columbidae (Col) | 4 | 3 | 1.6 | 1 | 1.6 | 1.3 | 2.5 | 4 | 2.5 |
| Corvidae (Cor) | 644 | 313 | 448.6 | 94.5 | 448.6 | 2.1 | 1.4 | 6.8 | 1.4 |
| Diomedeidae (Dio) | 53,013 | 7 | 0 | 0 | 0 | 7573.3 | ∞ | ∞ | ∞ |
| Estrildidae (Est) | 6 | 6 | 0 | 0 | 0 | 1 | ∞ | ∞ | ∞ |
| Falconidae (Fal) | 349 | 131 | 109.1 | 48 | 110.8 | 2.7 | 3.2 | 7.3 | 3.1 |
| Fregatidae (Fre) | 7,239 | 7,230 | 3 | 1 | 3 | 1 | 2413 | 7239 | 2413 |
| Fringillidae (Fri) | 3 | 0 | 0 | 0 | 0 | ∞ | ∞ | ∞ | ∞ |
| Gaviidae (Gav) | 33 | 3 | 13 | 5 | 13 | 11 | 2.5 | 6.6 | 2.5 |
| Gruidae (Gru) | 15 | 2 | 9 | 8 | 9 | 7.5 | 1.7 | 1.9 | 1.7 |
| Haematopodidae (Hae) | 24 | 5 | 1 | 1 | 1 | 4.8 | 24 | 24 | 24 |
| Hirundinidae (Hir) | 6 | 6 | 0 | 0 | 0 | 1 | ∞ | ∞ | ∞ |
| Hydrobatidae (Hyd) | 1 | 0 | 0 | 0 | 0 | ∞ | ∞ | ∞ | ∞ |
| Icteridae (Ict) | 84 | 31 | 14 | 13 | 14.1 | 2.7 | 6 | 6.5 | 6 |
|  | This study | WAHIS | GISAID | NCBI | GISAID+NCBI | WAHIS | GISAID | NCBI | GISAID+NCBI |
| Laridae (Lar) | 24,148 | 2,809 | 439 | 204.6 | 450.8 | 8.6 | 55 | 118 | 53.6 |
| Oceanitidae (Oce) | 1 | 1 | 0 | 0 | 0 | 1 | ∞ | ∞ | ∞ |
| Odontophoridae (Odo) | 40 | 1 | 0 | 0 | 0 | 40 | ∞ | ∞ | ∞ |
| Pandionidae (Pan) | 7 | 1 | 5 | 3 | 5 | 7 | 1.4 | 2.3 | 1.4 |
| Parulidae (Par) | 1 | 1 | 0 | 0 | 0 | 1 | ∞ | ∞ | ∞ |
| Passerellidae (Pas) | 3 | 3 | 0 | 0 | 0 | 1 | ∞ | ∞ | ∞ |
| Pelecanidae (Pel) | 55,947 | 1,901 | 161.6 | 127.8 | 176.3 | 29.4 | 346.2 | 437.8 | 317.3 |
| Phaethontidae (Phae) | 1 | 0 | 0 | 0 | 0 | ∞ | ∞ | ∞ | ∞ |
| Phalacrocoracidae (Phal) | 207,990 | 387 | 75.4 | 35 | 81 | 537.4 | 2758.5 | 5942.6 | 2567.8 |
| Phasianidae (Phas) | 69 | 5 | 17 | 14 | 17 | 13.8 | 4.1 | 4.9 | 4.1 |
| Phoenicopteridae (Phoe) | 262 | 34 | 0 | 0 | 0 | 7.7 | ∞ | ∞ | ∞ |
| Platylophidae (Pla) | 1 | 0 | 1 | 0 | 1 | ∞ | 1 | ∞ | 1 |
| Podicipedidae (Pod) | 278 | 43 | 60.1 | 25.9 | 60.9 | 6.5 | 4.6 | 10.7 | 4.6 |
| Procellariidae (Proce) | 543 | 43 | 24 | 3 | 24 | 12.6 | 22.6 | 181 | 22.6 |
| Psittacidae (Psici) | 173 | 61 | 3.8 | 1.8 | 3.8 | 2.8 | 45.5 | 96.1 | 45.5 |
| Psittaculidae (Psicu) | 1 | 0 | 0 | 0 | 0 | ∞ | ∞ | ∞ | ∞ |
| Rallidae (Ral) | 108 | 8 | 1 | 1 | 1 | 13.5 | 108 | 108 | 108 |
| Ramphastidae (Ram) | 2 | 2 | 0 | 0 | 0 | 1 | ∞ | ∞ | ∞ |
| Rheidae (Rhe) | 5 | 1 | 2 | 0 | 2 | 5 | 2.5 | ∞ | 2.5 |
| Scolopacidae (Sco) | 143 | 27 | 93.4 | 44.4 | 100 | 5.3 | 1.5 | 3.2 | 1.4 |
| Spheniscidae (Sph) | 3,798 | 435 | 2.9 | 3.8 | 3.8 | 8.7 | 1309.7 | 999.5 | 999.5 |
| Stercorariidae (Ste) | 61 | 50 | 2.5 | 2 | 2.6 | 1.2 | 24.4 | 30.5 | 23.5 |
| Strigidae (Stri) | 806 | 173 | 496.8 | 281.6 | 498.9 | 4.7 | 1.6 | 2.9 | 1.6 |
| Struthionidae (Stru) | 1 | 1 | 0 | 0 | 0 | 1 | ∞ | ∞ | ∞ |
| Sturnidae (Stu) | 2 | 2 | 0 | 0 | 0 | 1 | ∞ | ∞ | ∞ |
| Sulidae (Sul) | 197,827 | 777 | 108.8 | 18 | 115 | 254.6 | 1818.3 | 10990.4 | 1720.2 |
| Threskiornithidae (Thr) | 196 | 16 | 2 | 1 | 2 | 12.3 | 98 | 196 | 98 |
| Turdidae (Tur) | 2 | 1 | 2 | 2 | 2 | 2 | 1 | 1 | 1 |
| Tyrannidae (Tyr) | 1 | 1 | 0.9 | 0 | 0.9 | 1 | 1.1 | ∞ | 1.1 |
| Unidentified wild bird | 5,210 | 0 | 4 | 3 | 4 | ∞ | 1302.5 | 1736.7 | 1302.5 |
| **Total** | **615,883** | **16,902** | **6716.9** | **3772.5** | **7026.9** | **36.4** | **91.7** | **163.3** | **87.6** |
|  | This study | WAHIS | GISAID | NCBI | GISAID+NCBI | WAHIS | GISAID | NCBI | GISAID+NCBI |
| **Wild mammals** |  |  |  |  |  |  |  |  |  |
| Canidae (Can) | 166 | 164 | 118.1 | 18.3 | 118.1 | 1 | 1.4 | 9.1 | 1.4 |
| Castoridae (Cast) | 2 | 0 | 0 | 0 | 0 | ∞ | ∞ | ∞ | ∞ |
| Cricetidae (Cri) | 16 | 14 | 0 | 0 | 0 | 1.1 | ∞ | ∞ | ∞ |
| Delphinidae (Del) | 82 | 2 | 9 | 7.1 | 9.1 | 41 | 9.1 | 11.5 | 9 |
| Didelphidae (Did) | 7 | 7 | 2.1 | 0 | 2.1 | 1 | 3.3 | ∞ | 3.3 |
| Felidae (Fel) | 36 | 36 | 12.3 | 3 | 12.3 | 1 | 2.9 | 12 | 2.9 |
| Leporidae (Lep) | 1 | 1 | 0 | 0 | 0 | 1 | ∞ | ∞ | ∞ |
| Mephitidae (Mep) | 139 | 137 | 105 | 12.9 | 106 | 1 | 1.3 | 10.8 | 1.3 |
| Mustelidae (Mus) | 55 | 11 | 5.9 | 2.1 | 5.9 | 5 | 9.3 | 26.2 | 9.3 |
| Otariidae (Ota) | 35,424 | 3,274 | 33.5 | 32.3 | 36.8 | 10.8 | 1057.4 | 1096.7 | 962.6 |
| Phocidae (Phoci) | 27,375 | 2,620 | 51.1 | 17.1 | 51.1 | 10.4 | 535.7 | 1600.9 | 535.7 |
| Phocoenidae (Phoco) | 40 | 0 | 3 | 3 | 3 | ∞ | 13.3 | 13.3 | 13.3 |
| Procyonidae (Procy) | 48 | 45 | 20.9 | 10 | 22 | 1.1 | 2.3 | 4.8 | 2.2 |
| Sciuridae (Sci) | 6 | 1 | 0 | 0 | 0 | 6 | ∞ | ∞ | ∞ |
| Ursidae (Urs) | 12 | 11 | 4.9 | 2.9 | 4.9 | 1.1 | 2.4 | 4.1 | 2.4 |
| **Total** | **63,409** | **6,323** | **365.8** | **108.6** | **371.3** | **10** | **173.3** | **583.9** | **170.8** |

**Fig 1.**
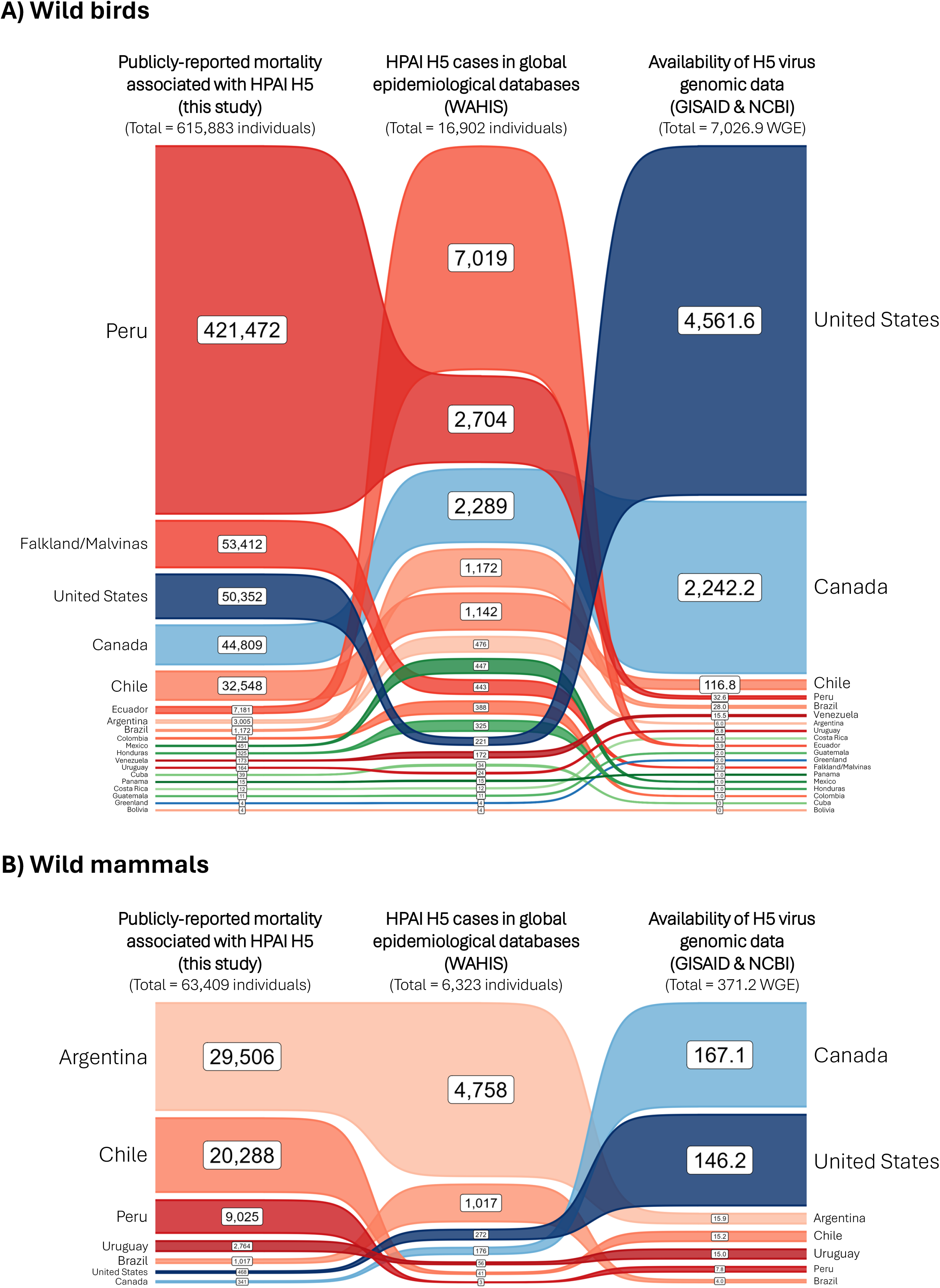
Comparison between publicly-reported wildlife mortality associated with HPAI H5 (this study), wildlife reportedly affected by HPAI H5 in global epidemiological databases (WAHIS), and the availability of H5 virus genomic data (GISAID & NCBI) for wild birds (A) and wild mammals (B) among countries/territories in the Americas. Band thickness is proportional to the percentage of the column total, and bands are ordered by rank (descending). Countries/territories are colored by continent (Northern America = blue, Central America/Caribbean = green, South America = red). Genomic data is quantified in whole genome equivalents (WGE; number of gene segment sequences divided by eight). Study period: November 2021 – July 2024 (inclusive).

**Fig 2.**
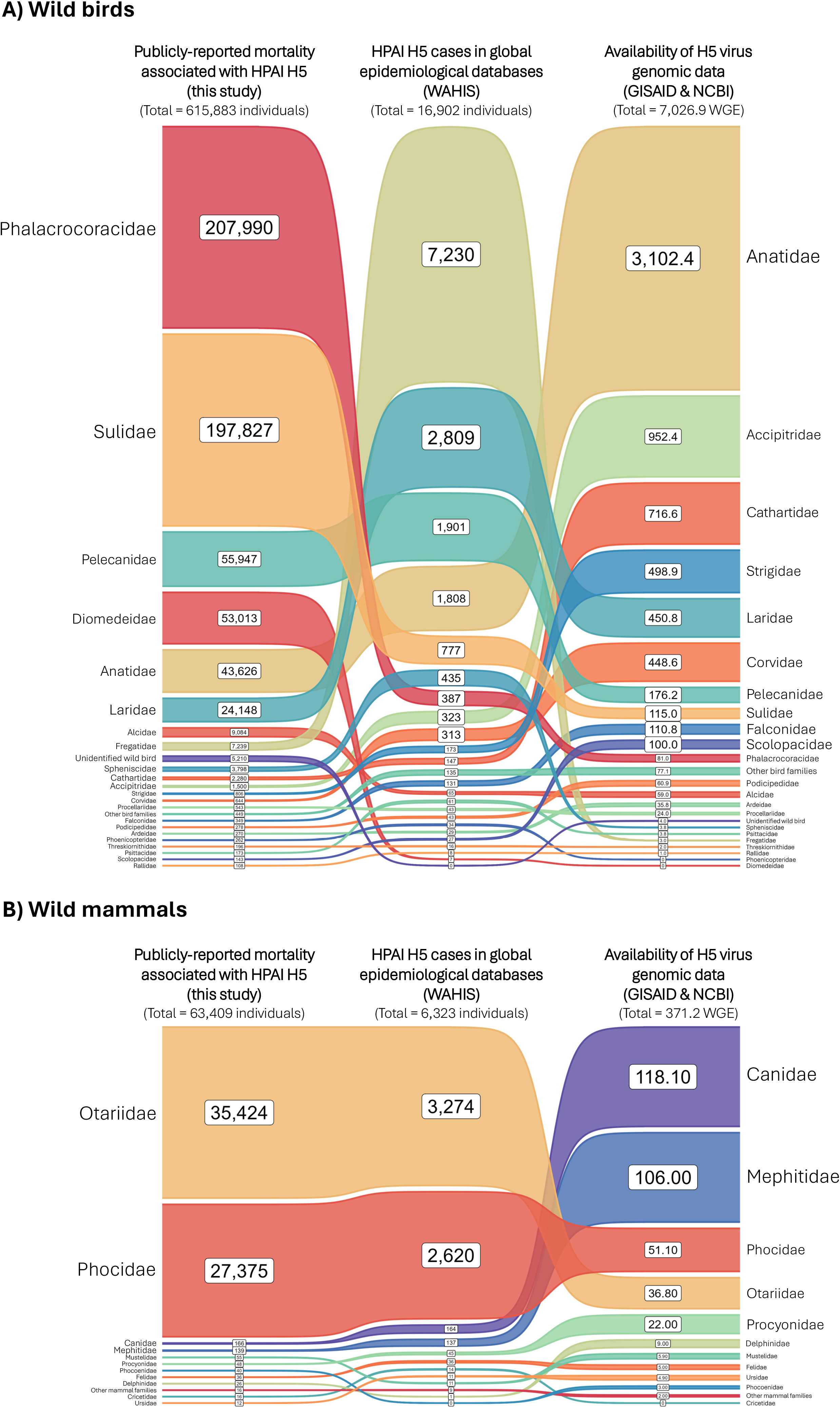
Comparison between publicly-reported wildlife mortality associated with HPAI H5 (this study), wildlife reportedly affected by HPAI H5 in global epidemiological databases (WAHIS), and the availability of H5 virus genomic data (GISAID & NCBI) among host families of wild birds (A) and wild mammals (B) in the Americas. Band thickness is proportional to the percentage of the column total, and bands are ordered by rank (descending). Genomic data is quantified in whole genome equivalents (WGE; number of gene segment sequences divided by eight). Study period: November 2021 – July 2024 (inclusive).

### Wildlife affected by HPAI H5 in epidemiological databases (WAHIS and EMPRES-i+)

WAHIS-reported wildlife affected by HPAI H5 in the Americas comprised 16,902 wild birds (261 species, 45 families) and 6,323 wild mammals (31 species, 13 families). Relative to the consolidated dataset, WAHIS had representation ratios of 1:36 for wild birds and 1:10 for wild mammals (i.e. for each 1 wild bird or wild mammal affected by HPAI H5 in WAHIS there were 36 wild bird and 10 wild mammal deaths associated with HPAI H5 in our consolidated dataset) (Tables 1 and 2).

The extent to which publicly-reported mortality was represented in WAHIS data was noticeably uneven among countries/territories (Fig 1). For instance, while Peru had the highest total estimated wild bird mortality (ranking 1^st^ with 68.4% of publicly-reported wild bird mortality), it ranked 2^nd^ in WAHIS-reported affected wild birds (representing 16.0% of wildlife affected by HPAI H5 reported to WAHIS). Similarly, while the USA ranked 3^rd^ in publicly-reported wild bird mortality (8.2%), it was only 11^th^ in WAHIS-reported affected wild birds (1.3%). In contrast, although Ecuador ranked 6^th^ in publicly-reported wild bird mortality (1.2%), it was 1^st^ in WAHIS-reported affected wild birds (41.5%). A similar pattern was observed for wild mammals, as exemplified by Chile ranking 2^nd^ in publicly-reported wild mammal mortality (32.0%) but only 6^th^ in WAHIS-reported affected wild mammals (0.6%), whereas Brazil ranked 5^th^ in publicly-reported wild mammal mortality (1.6%) but was 2^nd^ in WAHIS-reported affected wild mammals (16.1%). In terms of representation in WAHIS versus publicly-reported mortality, the countries/territories with the lowest relative representation were United States (1:228), Peru (1:156), and the Falkland/Malvinas Islands (1:121) for wild birds and Peru (1:3008), Chile (1:495), and Uruguay (1:49) for wild mammals (Table 1 and Fig 3A).

**Fig 3.**
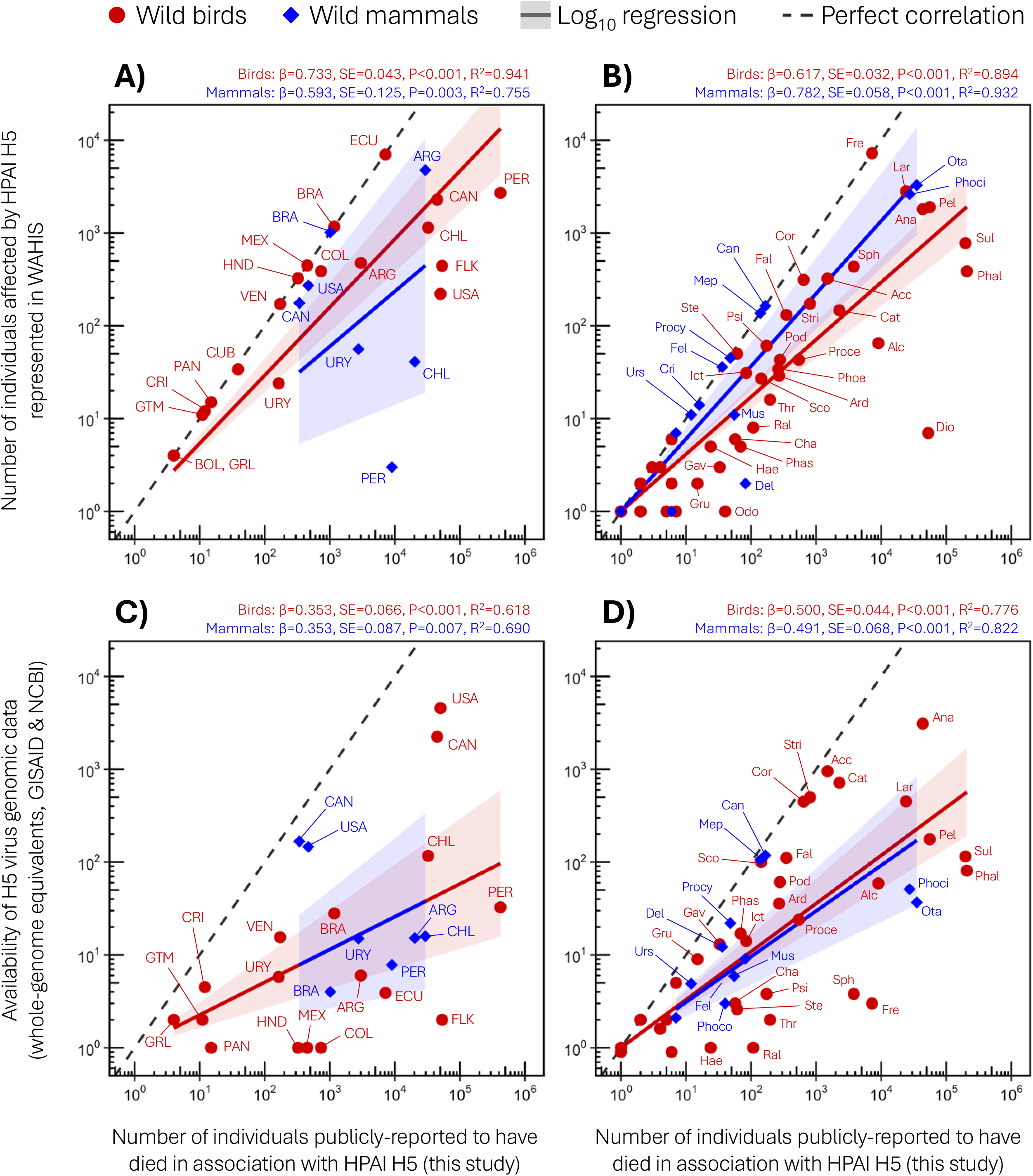
Correlation between publicly-reported wildlife mortality associated with HPAI H5 and WAHIS-reported affected wildlife (A, B) or the availability of H5 virus genomic data from wildlife (C, D) in the Americas. Countries/territories (A, C) and host families (B, D) are labelled as in Tables 1 and 2. Host families <10 publicly-reported deaths are not labelled. Regression lines (and their 95% confidence interval in shaded areas) are drawn separately for wild birds (red) and wild mammals (blue) and regression parameters are shown above each plot. Study period: November 2021 – July 2024 (inclusive).

These country-level differences also translated into discrepancies in how host families were represented in WAHIS data (Figs 2 and 3B). For instance, while the families Phalacrocoracidae and Sulidae ranked 1^st^ and 2^nd^ in publicly-reported wild bird mortality (33.8% and 32.1%, respectively), they ranked 7^th^ and 5^th^ in WAHIS-reported affected wild birds (2.3% and 4.6%, respectively). In contrast, families Fregatidae and Laridae ranked 8^th^ and 6^th^ in publicly-reported wild bird mortality (1.2% and 3.9%, respectively), yet were 1^st^ and 2^nd^ in WAHIS-reported affected wild birds (42.8% and 16.6%, respectively). On the other hand, although wild mammals were generally underrepresented in WAHIS data, the ranking order of host families was generally consistent with that in publicly-reported mortality (Fig 2B). In terms of overall relative representation, the host families with the lowest relative representation were Diomedeidae (1:7573), Phalacrocoracidae (1:537), and Sulidae (1:255) for wild birds and Delphinidae (1:41), Otariidae (1:11), and Phocidae (1:10) for wild mammals (Table 2 and Fig 3B). At the species level, WAHIS data represented 90.1% (261/287) of wild bird species and 79.5% (31/39) of wild mammal species for which there was publicly-reported mortality associated with HPAI H5, and this representativeness was generally consistent both across countries/territories (S1 Table and S1 Fig) and host families (S2 Table and S1 Fig).

Because EMPRES-i+ does not provide the number of individuals affected in each disease event, we assessed its representation at the species level. EMPRES-i+ data represented a limited host diversity for HPAI H5 detections in wild birds (105 species, 28 families) and wild mammals (27 species, 13 families). Compared to our consolidated mortality dataset, EMPRES-i+ represented only 36.6% (105/287) of wild bird species and 69.2% (27/39) of wild mammal species involved in HPAI H5 events, and this species-level coverage varied considerably across both countries/territories (S1 Table and S1 Fig) and host families (S2 Table and S1 Fig). Additionally, when crosschecking information with other sources it became clear that EMPRES-i+ data had some relevant inconsistencies, such as the incorrect attribution of HPAI H5 cases to sea otter (*Enhydra lutris*) instead of marine otter (*Lontra felina*) (event ID 347347), red-breasted goose (*Branta ruficollis*) instead of red-breasted merganser (*Mergus serrator*) (event ID 316129), grey petrel (*Procellaria cinerea*) instead of sooty shearwater (*Ardenna grisea*) (event ID 382498), among others, likely arising from faulty automated processing or human error while importing data from WAHIS to EMPRES-i+.

### HPAI H5 wildlife virus data in genomic databases (GISAID and NCBI)

Considered separately, GISAID and NCBI data respectively comprised 6,716.9 WGE (whole genome equivalents; i.e. number of gene segment sequences divided by eight) and 3,772.5 WGE for wild birds, and 365.8 WGE and 108.6 WGE for wild mammals in the Americas (Tables 1 and 2). The merged GISAID/NCBI dataset (S4 File) amounted to 7,026.9 WGE for wild birds (175 species, 36 families) and 371.3 WGE for wild mammals (26 species, 11 families). Relative to the consolidated mortality dataset, the representation ratio of GISAID/NCBI data was 1:88 for wild birds and 1:171 for wild mammals.

The availability of H5 virus genomic data was markedly heterogeneous across countries (Fig 1 and Table 1). Available H5 virus genomic data from wild birds predominantly originated from the United States (4,561.6 WGE; 64.9%) and Canada (2,242.3 WGE; 31.9%), and for wild mammals this pattern was reversed with Canada (167.1 WGE; 45.0%) followed by the United States (146.3 WGE; 39.4%). As a result, genomic representation of publicly-reported wildlife mortality differed sharply between the United States and Canada versus the rest of the Americas. For the United States and Canada combined, representation ratios were 1:14 for wild birds and 1:2.6 for wild mammals, whereas for all other American countries combined the ratios were 1:2335.1 for wild birds and 1:1081.6 for wild mammals, respectively. Specifically, the countries/territories with the lowest genomic representation were the Falkland/Malvinas Islands (1:26706), Peru (1:12919), and Ecuador (1:1853) for wild birds, and Argentina (1:1859), Chile (1:1330), and Peru (1:1165) for wild mammals (Table 1 and Fig 3C).

These country-level differences resulted in asymmetries in H5 virus genomic data availability among host families (Figs 2 and 3D). While families Phalacrocoracidae and Sulidae respectively ranked 1^st^ and 2^nd^ in publicly-reported wild bird mortality, they ranked 11^th^ and 8^th^ in genomic data availability with representation ratios of 1:2568 and 1:1720, respectively. In contrast, families Anatidae and Accipitridae ranked 5^th^ and 12^th^ in publicly-reported wildlife mortality, yet they ranked 1^st^ and 2^nd^ in genomic data availability with representation ratios of 1:14 and 1:1.6, respectively. Similar discrepancies were also noted for mammals. Otariidae and Phocidae ranked 1^st^ and 2^nd^ in publicly-reported wild mammal mortality, but ranked 4^th^ and 3^rd^ in genomic data availability with relative representation of 1:963 and 1:536, respectively. In contrast, families Canidae and Mephitidae ranked 3^rd^ and 4^th^ in publicly-reported wild mammal mortality, but ranked 1^st^ and 2^nd^ in genomic data availability with relative representation of 1:1.4 and 1:1.3, respectively. At the species level, genomic data represented 61.0% (175/287) of wild bird species and 66.7% (26/39) of wild mammal species for which there was publicly-reported mortality associated with HPAI H5, and this representativeness varied considerably across both countries/territories (S1 Table and S1 Fig) and host families (S2 Table and S1 Fig).

## Discussion

This study represents the most comprehensive compilation of public information on wildlife mortality associated with HPAI H5 during the first wave of dissemination in the Americas, up to July 2024. Because the dataset relies on reported information, it is necessarily affected by variation in observation probability (e.g. wildlife mortality at sea or in remote areas is less likely to be observed), access to carcasses (e.g. observers may lack permission or access to document wildlife mortalities), reporting incentives (e.g. observers may consider wildlife mortality unremarkable or undesirable to be publicized or may lack access or permission to report it), diagnostic confirmation (e.g. testing of poultry or livestock samples may have been prioritized over wildlife samples in some jurisdictions or stages of an outbreak), and publication delays (e.g. timely publication of some reports on wildlife mortalities may have been hindered by contingencies or limited resources). Thus, while the dataset represents the best-feasible compilation of public information to date, it should not be interpreted as a definitive mortality census, as it may have underestimated wildlife mortality in instances where carcasses were not detected, counted, tested, or publicly reported. Conversely, because the dataset includes all wildlife mortality associated with HPAI H5 detections, regardless of molecular or pathological confirmation of individual cases, some deaths may have been coincidentally associated with HPAI H5 but caused by other factors. This may be particularly relevant in Peru and northern Chile, where HPAI H5 outbreaks in 2023/2024 overlapped with a strong El Niño event that may have contributed to the mortality of marine wildlife, which is a well-documented cause of seabird mass starvation and nest desertion in the region [41–43]; nevertheless, there is robust evidence that HPAI H5 played a major role driving wildlife mortality in this region at the time [44–48]. These limitations are important, but they do not undermine the central comparison of this study, where we assess whether international databases provide a representative and interpretable framework for public information on wildlife mortality in the Americas associated with HPAI H5.

Our results show that 615,883 wild birds and 63,409 wild mammals were publicly-reported to have been found dead or estimated to have died in association with HPAI H5, with the vast majority reported from South America (84.4% of wild birds and 98.7% of wild mammals). While these numbers are small compared to losses in the poultry industry –the United States alone reported the loss of >175 million domestic birds over the same period [49]–, they are acutely concerning because many affected wildlife species have far smaller population sizes and slower recovery potential. For most wildlife species, the lack of pre-outbreak population estimates makes it difficult to quantify conservation impacts. Yet, for the few species where pre- and post-outbreak population data are available, the impacts of HPAI H5 have been dramatic, especially for colonial marine species: ∼28% population decrease for South American sea lions (*Otaria flavescens*) in northern Argentine Patagonia [50], ∼35% breeding population decrease for Peruvian pelicans (*Pelecanus thagus*) in marine protected areas in Peru [48], ∼60% breeding population decrease for Humboldt penguins (*Spheniscus humboldti*) in northern and central Chile [51], and ∼95% pup mortality and ∼45% adult population decrease for southern elephant seals (*Mirounga leonina*) in Argentina [52,53]. These losses are compounded by additional ecological pressures such as fisheries competition and bycatch, climate change, and marine pollution, among others [54,55], which may jeopardize their resilience and ability to recover from the impacts of HPAI H5.

These examples illustrate why wildlife mortality data are not only epidemiological records but also essential conservation evidence. If mortality events are underrepresented or unevenly represented in global systems, conservation risk may be underestimated for species, regions or ecosystems with limited reporting or sequencing capacity. Moreover, wildlife also plays a significant role in HPAI H5 ecology and evolution. This is illustrated by the recent emergence of reassortant HPAI H5 strains that combine genomic segments from viruses of Eurasian Goose/Guangdong lineage with those from low pathogenicity strains circulating in wild birds in North America [36,56,57] and South America [58,59]. Furthermore, the persistent circulation of HPAI H5 in wild mammals can lead to the emergence of viral strains carrying mammalian-adaptive mutations, with implications for zoonotic risk [17,18,39]. Accurate interpretation of wildlife data is therefore important for conservation, animal health, and public health.

The discrepancies documented here likely reflect a combination of biological, logistical, institutional, and socioeconomic factors. Wildlife mortality is difficult to detect in remote areas or at sea; diagnostic and sequencing capacity varies among countries; reporting requirements and practices differ among authorities and sectors; and wildlife samples may compete with domestic animal samples for limited laboratory resources. Differences in WOAH reporting practices among countries can also cause heterogeneity in how different countries and species are represented in WAHIS. Some countries have reported nearly every confirmed or suspected HPAI H5 case in wildlife (e.g. Brazil, Ecuador), while others only reported cases that were government laboratory-confirmed (e.g. Chile), or reported the first laboratory-confirmed case for each host species but then did not consistently report additional suspected or confirmed cases (e.g. United States). In jurisdictions where only governmental authorities are allowed or expected to share information on wildlife mortality or suspected HPAI H5 cases in any public space, WAHIS may become the only public source of information. In such settings, apparent agreement between WAHIS and other public sources may reflect limited public reporting pathways rather than a complete picture of wildlife mortality.

It is not unexpected that large, collaborative, multinational epidemiological databases may have coverage gaps, reporting heterogeneity, and occasional data inconsistencies [40,60]. However, the magnitude and directionality of the gaps documented here have important implications for the interpretation of studies that rely on these resources. The issue is therefore not that databases such as WAHIS or EMPRES-i+ lack value; rather, our results show that their records must be interpreted cautiously in light of the surveillance, reporting, and data-entry contexts that produced them. For instance, analyses based on WAHIS or EMPRES-i+ data to summarize the epidemiological patterns of HPAI H5 in South America (e.g. [23,26]) may capture not only the distribution of HPAI H5 in wildlife, but also differences in surveillance capacity and intensity, reporting thresholds, case definitions, and data-entry and submission procedures among countries and host taxa. As a result, apparent geographic, temporal or taxonomic patterns may partly reflect how data were generated and reported rather than where wildlife mortality or viral circulation has been greatest.

This has implications for studies relying on international databases to summarize HPAI H5 distribution or inter-annual patterns (e.g. [22]), assess zoonotic risks (e.g. [27]), infer host susceptibility (e.g. [25]), model ecological predictors (e.g. [28]), or assess conservation impacts. Simply including a statement acknowledging that public databases are incomplete may be insufficient when data gaps are large, uneven, and structured by reporting or sequencing effort. For quantitative epidemiological, ecological, or conservation studies, database records should ideally be crosschecked and validated against national reports, scientific publications, field-survey data, and other complementary sources. Where validation is not possible, analyses should explicitly address how reporting gaps could affect interpretation, for example through sensitivity analysis, stratification by reporting coverage, or cautious framing of conclusions. Likewise, animal health authorities, wildlife managers and other decision-makers should avoid treating global databases as complete mortality datasets and should interpret them considering the surveillance and reporting context in which they were produced.

Our results also demonstrate profound discrepancies between countries and taxa with the greatest availability of H5 virus genomic data from wildlife versus those with the highest publicly-reported wildlife mortality. These discrepancies likely reflect differences in laboratory capacity, sequencing resources, sample quality, prioritization, and timeliness of public sequence sharing. Virus genome sequencing can be expensive and is not mandatory for diagnostic confirmation [61]. In resource-constrained settings, national reference laboratories may prioritize diagnostic testing over sequencing, and samples from poultry or domestic animals may be prioritized for sequencing over those from wildlife. Timeliness is also important. Our analysis used GISAID/NCBI data available one year after the end of study period, yet it is not uncommon for H5 virus genomic data to be published >400 days after sample collection [33]. For instance, the Falkland/Malvinas Islands had limited publicly-available H5 virus genomic data in this study (only 2 WGE despite >50,000 wild bird deaths), but preprint articles indicate that substantial genomic datasets should become available in the near future [62,63]. Such delays limit the usefulness of public genomic data for real-time risk assessment and decision-making during unfolding epizootics [33].

The taxonomic distribution of genomic data also requires careful interpretation. The top four wild bird families with greatest H5 virus genomic data availability –Anatidae, Accipitridae, Cathartidae, and Strigidae– had the bulk of their publicly-reported mortality (>90%) in the United States and Canada. Together, these families comprised 75% of the H5 virus genomic data from wild birds but <10% of publicly-reported wild bird mortality in the Americas, corroborating that international differences in sequencing effort, rather than epidemiological patterns, are the main driver of genomic data availability. Other factors may also influence sequencing patterns, including prioritization of species that are threatened and/or intensively managed (e.g. California condor *Gymnogyps californicus* with a representation ratio of 1:2.0), charismatic and/or culturally-symbolic (e.g. bald eagle *Haliaeetus leucocephalus* with a representation ratio of 1:1.8), or of special interest for the surveillance of emerging mammal-adapted strains of zoonotic concern (e.g. red fox *Vulpes vulpes* with a representation ratio of 1:1.4). Genomic availability may also reflect targeted surveillance and sequencing efforts in hosts that are classically recognized as playing a role in the ecology and long-distance dissemination of avian influenza viruses (e.g. family Scolopacidae with a representation ratio of 1:1.3). Furthermore, sampling strategies that rely on hunter-harvested specimens may also result in some host families having a higher proportion of freshly-collected samples, which can translate into better RNA integrity and result in greater sequencing success (e.g. family Anatidae with a representation ratio of 1:13.4). Another factor is that during localized large-scale outbreaks only a subset of individuals will be sampled for virus genome sequencing. Consequently, species for which mortality was predominantly clustered in large-scale outbreaks (e.g. Caspian tern *Hydropogne caspia* and northern gannet *Morus bassanus* with representation ratios of 1:127.7 and 1:284.8, respectively) may be less represented in H5 virus genomic data on a per-individual basis than species with sporadic cases across several jurisdictions (e.g. black vulture *Coragyps atratus* and great horned owl *Bubo virginianus* with representation ratios of 1:3.8 and 1:1.2, respectively).

These blind spots matter because genomic representation should not be interpreted as indicative of higher viral circulation (e.g. [65]) or epidemiological relevance. For example, HPAI H5 viruses from pinnipeds in South America are scarcely represented in genomic databases (<45 WGE) despite being associated with >50,000 deaths across five countries and with robust evidence of mammal-mammal transmission [17,18,66]. In contrast, HPAI H5 viruses from foxes and skunks in North America are far better represented in genomic data (>220 WGE) despite being involved in only ∼300 deaths in two countries, without evidence of sustained mammal-mammal transmission [8,12]. Studies and applications that use genomic databases to assess zoonotic risk, monitor mutations of public health concern, design vaccines or diagnostics, or infer viral spatiotemporal dissemination should therefore account for regional and taxonomic differences in sequencing effort.

Several practical measures could make international HPAI H5 data more useful for wildlife health and conservation decisions. Detailed recommendations are summarized in Table 3. Briefly, epidemiological databases could allow for data entry that distinguishes between laboratory-confirmed cases, field-counted mortality, estimated mortality from surveys, and population-level mortality estimates; include uncertainty or confidence levels; use standardized taxonomic identifiers; and document whether records represent index cases, partial updates, or complete event summaries. Automated data transfers between platforms should include human-supervised validation to reduce taxonomic and geographic errors. Genomic databases could improve utility by encouraging timely sequence submission, clearer linkage between sequence records and epidemiological events, and metadata describing sampling strategy. Finally, international agencies and funders can strengthen HPAI H5 interpretation by supporting wildlife-specific surveillance, diagnostic, sequencing, and data-management capacity, particularly in regions where wildlife mortality is high but laboratory resources are limited.

**Table 3.**
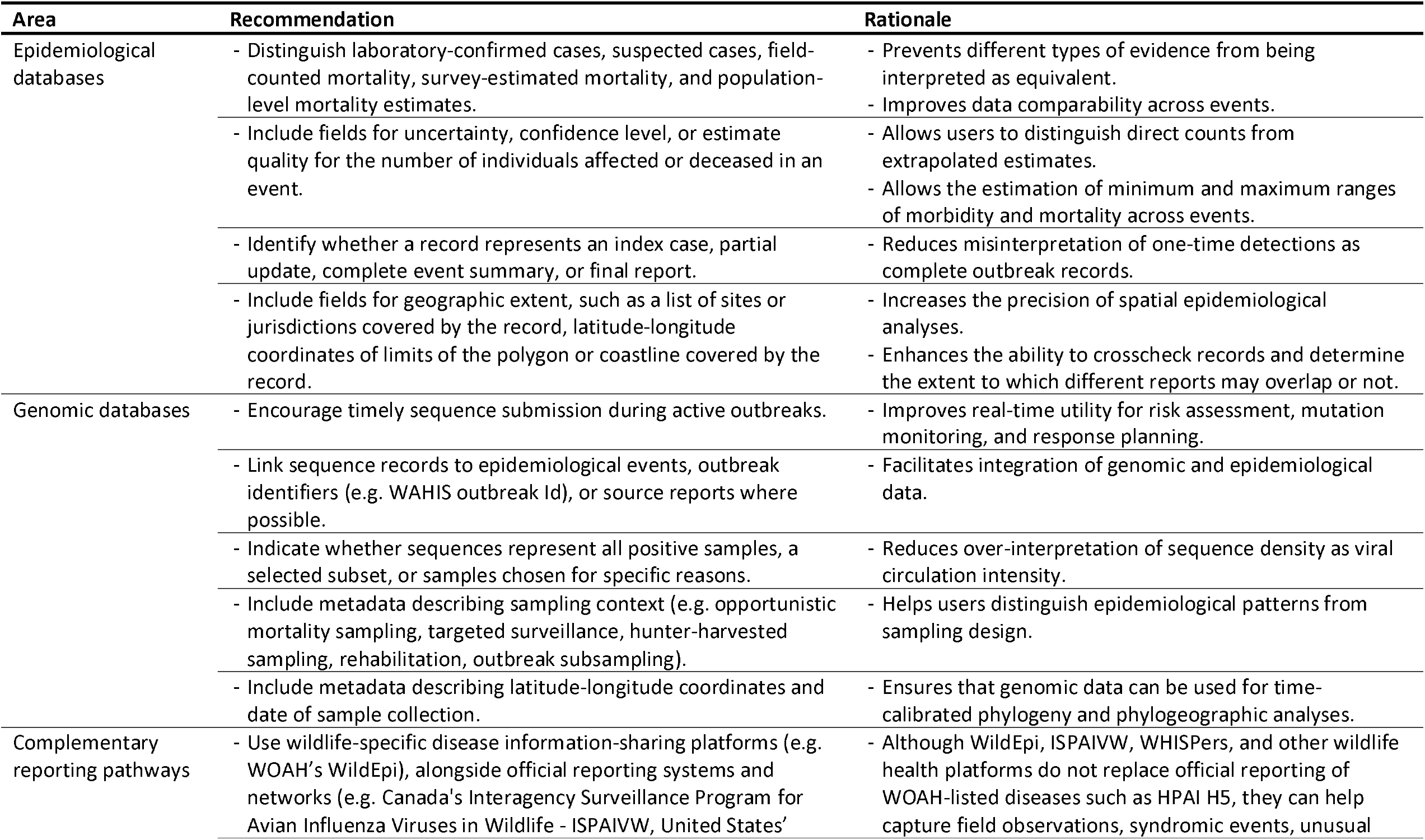
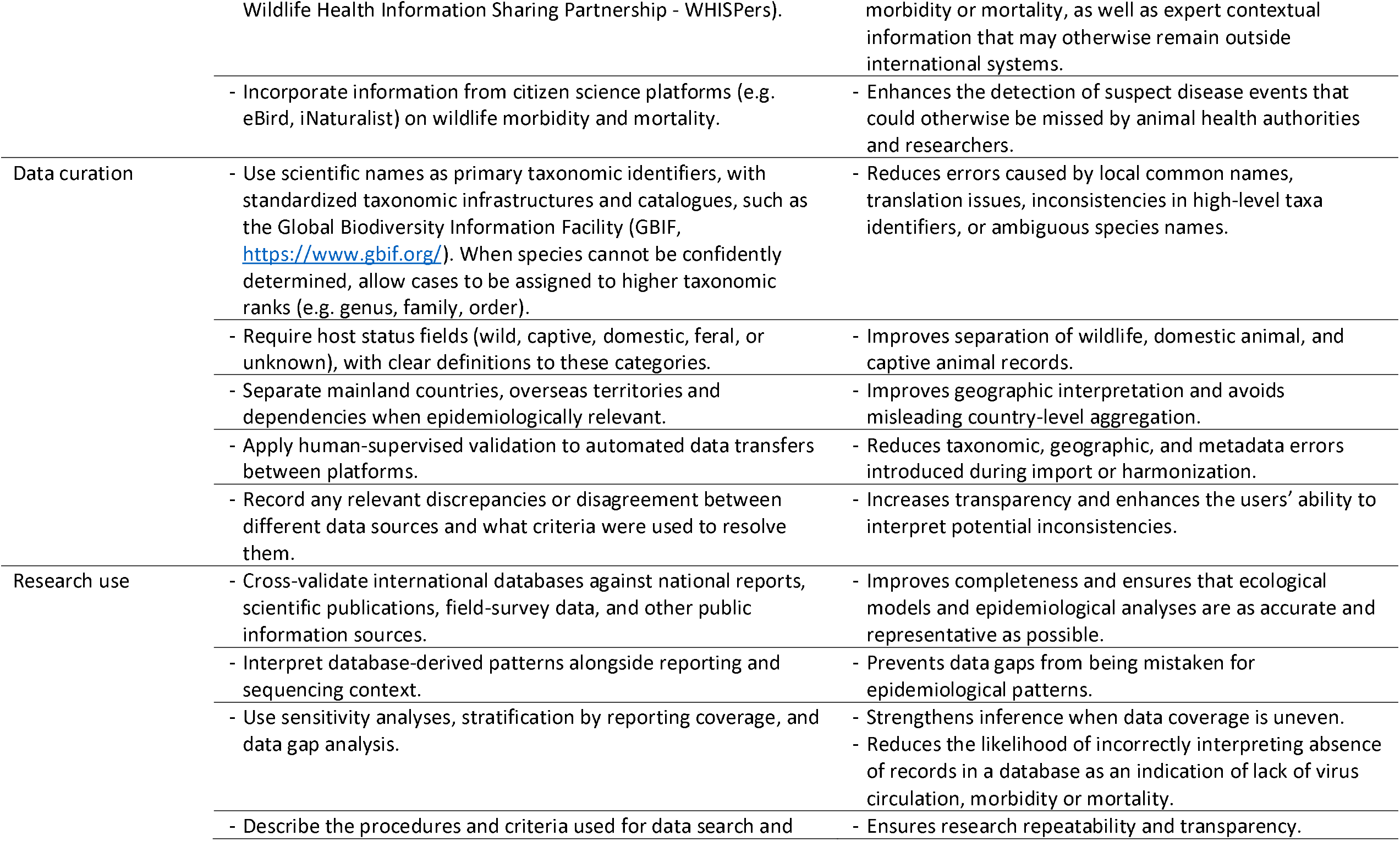
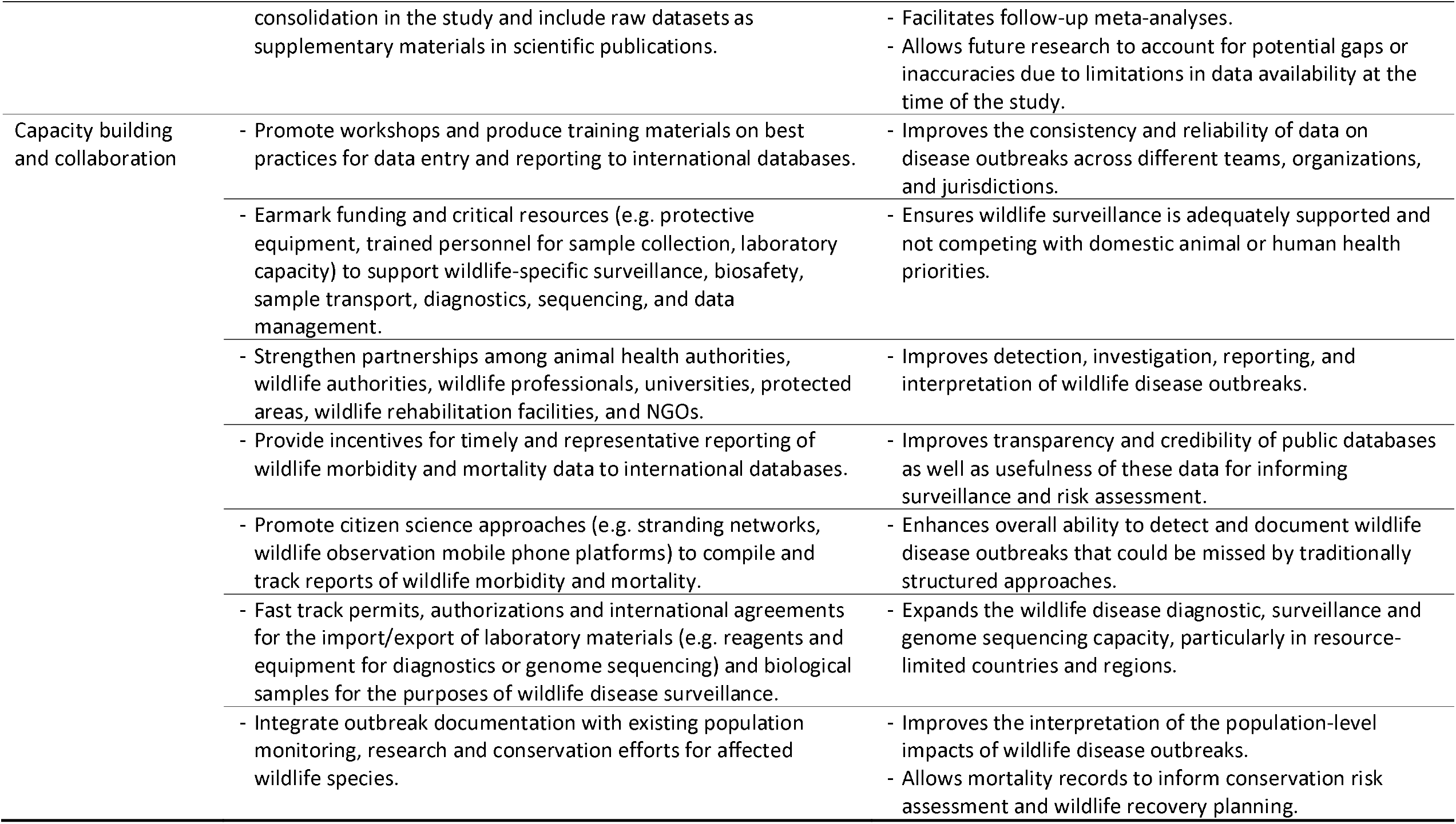
Practical recommendations to improve the interpretation and value for One Health surveillance, risk assessment and conservation action of international HPAI H5 wildlife data.

More broadly, the discrepancies in HPAI H5 surveillance, reporting, and genomic data availability in wildlife are largely driven by human factors such as socioeconomic, institutional, and political realities. Strengthening field surveillance and laboratory capacity through collaboration among national animal health agencies, wildlife managers, wildlife biologists and veterinarians, universities, protected area authorities, and non-governmental organizations would improve the investigation and reporting of HPAI and other infectious diseases in wildlife worldwide. Emerging wildlife-specific information systems may help address some of these broader challenges. For example, WOAH’s upcoming WildEpi system will be an information-sharing platform for non-listed wildlife health events, complementing official reporting through WAHIS [67]. Although HPAI H5 remains a WOAH-listed disease and therefore requires reporting through established mechanisms, WildEpi illustrates an important direction for wildlife health surveillance by creating dedicated pathways to capture field observations, syndromic events, unusual morbidity or mortality, and expert contextual information that may otherwise remain outside international systems. Such approaches could strengthen early detection and improve the flow of field-based information needed to interpret wildlife mortality events. Under a One Health paradigm, pathogen surveillance should encompass and appreciate the intrinsic, cultural, ecological, and economic value of wildlife, rather than merely considering wildlife as a potential vector or reservoir of risk to poultry, livestock, or humans. Earmarking resources specifically for wildlife surveillance, diagnostics, sequencing, data management, and safe field response would help ensure that wildlife needs do not have to compete entirely with domestic animal priorities in resource-constrained settings [68].

Lastly, it is worth noting that although this study focused on the Americas, there is reason to suspect that inconsistent reporting of wildlife mortality associated with HPAI is a global challenge. For instance, several notable wildlife mortalities associated with HPAI H5 in Eurasia during the study period appear to be incompletely represented in WAHIS, including 3,500 red knots (*Calidris canutus*) in Germany (October 2020 to March 2021) [69], 300 demoiselle cranes (*Grus virgo*) in India (November 2021) [70,71], 2,286 Dalmatian pelicans (*Pelecanus crispus*) in Greece (February to April 2022) [4], 20,531 sandwich terns (*Thalasseus sandvicensis*) across several countries in western Europe (April to June 2022) [6], 5,035 northern gannets (*Morus bassanus*) at Bass Rock in Scotland (June 2022) [5], and 3,500 northern fur seals (*Calorhinus ursinus*) in Russia (July 2023) [72]. As of July 2025, WAHIS data did not include reported HPAI H5 cases for demoiselle cranes or northern fur seals, and the numbers of individuals reported in WAHIS represent only a fraction of published mortality estimates for red knots (no individuals reported for Germany, 79 individuals reported for all of Europe), Dalmatian pelicans (600 individuals reported for Greece), sandwich terns (2,819 individuals reported for all of Europe), and northern gannets (199 individuals reported for the entire United Kingdom). These examples suggest that the challenges identified in the Americas are likely relevant to global HPAI H5 surveillance, wildlife mortality reporting, and conservation planning.

In conclusion, public epidemiological and genomic databases are invaluable resources for understanding wildlife infectious diseases such as HPAI H5. Our findings show that their value can be strengthened by interpreting database records alongside the surveillance, reporting, and sequencing contexts that produced them. For wildlife, where mortality detection is often opportunistic and conservation implications may be severe, integrating international databases with national reports, scientific publications, field observations, and genomic data is essential. Such an integrated approach can reduce the risk of mistaking gaps in data coverage for real-world epidemiological patterns, and can better support One Health surveillance, risk assessment, and conservation action.

## Materials and methods

### Scope definition

#### Geography

The study focused exclusively on the Americas, including the Caribbean, Greenland and oceanic islands closely associated with the continent (e.g. Falkland/Malvinas Islands, Galapagos Islands, Isla de Cocos). Bouvet Island, South Georgia, and the South Sandwich Islands were excluded. Country/territory names followed the United Nations’ M49 Standard [73].

#### Period

The study comprises cases from November 2021 to July 2024, inclusive. The rationale for this interval is that the first HPAI H5 case in the Americas was recorded on 26 November 2021 and the period of July-August 2024 marked a natural valley in the number of HPAI cases detected in the Americas [74,75].

#### Viruses

The study includes all strains of the influenza A virus (*Alphainfluenzavirus influenzae*) attributed to hemagglutinin subtype H5, except when sequence analysis established that the hemagglutinin gene was not derived from the Goose/Guangdong lineage.

#### Hosts

The study focuses on wild birds and wild mammals. The following taxa were excluded because they are not native to the Americas, are most often kept as pets or livestock (i.e. not zoological or conservation-oriented collections), or are invasive synanthropic species: *Anser anser* (greylag goose), *Bos taurus* (cattle), *Canis familiaris* (dog), *Capra hircus* (goat), *Columba livia* (rock pigeon), *Coturnix coturnix* (common quail), *Felis catus* (cat), *Mus musculus* (house mouse), Numididae (guineafowl), *Passer domesticus* (house sparrow), *Pavo cristatus* (common peafowl), *Phasianus colchicus* (common pheasant), *Sus scrofa* (pig), and *Vicugna pacos* (alpaca). Reports of mortality or HPAI H5 detection in *Anas platyrhynchos* (mallard) and *Cairina moschata* (muscovy duck) were excluded because although they could be native to some of the studied countries, it was not possible to reliably differentiate whether individual reports referred to wild or domestic individuals. Reports of mortality or HPAI H5 detection in *Meleagris gallopavo* (turkey) were only included for countries where the species is native (i.e. Canada and United States) and when the report explicitly referred to “wild turkey”. Reports assigned to non-specific taxa that may include domestic or invasive synanthropic species (“avian”, “Anatidae”, “duck”, “feline”, “goose”, “multispecies”, “peacock”, “peafowl”, “pheasant”, “pigeon”, “unknown”, and “waterfowl”) were excluded unless there was an explicit indication that the record referred to wildlife (e.g. the term “wild”) or there was another source indicating a confirmed wildlife case with matching metadata (location and collection date). Reports on HPAI H5 detections in humans, environmental samples, and laboratory-derived strains were excluded.

### Data aquisition and processing

Data were acquired on 31 July 2025, i.e. exactly one year after the end of the study period. WAHIS data were obtained both by submitting a user support request (https://wahis.woah.org/) and through consultation of the Events Management interface (https://wahis.woah.org/#/event-management); search criteria were set to include all H5 serotypes for the disease “Influenza A viruses of high pathogenicity (Inf. with) (non-poultry including wild birds) (2017-)”. EMPRES-i+ data were downloaded via the Epidemiology interface (https://empres-i.apps.fao.org/epidemiology); search criteria were set to include all events of “Influenza - Avian” with any H5 serotypes. GISAID data were downloaded via the EpiFlu interface (https://platform.epicov.org/epi3/); search criteria were set to include all type A subtype H5 isolates. NCBI data was obtained via the Influenza Virus Data Hub interface (https://www.ncbi.nlm.nih.gov/labs/virus/vssi/#/virus); search criteria were set to include all “Alphainfluenzavirus (taxid:197911)” of the genotype “H5*” isolates.

Further details on data processing are provided in S1 File. Briefly, each dataset was manually revised to exclude geographical areas, collection dates, and host taxa outside of the scope definition. For WAHIS, the number of affected individuals in each disease event was derived from the number of susceptible, cases, deaths and killed/disposed individuals. GISAID and NCBI data were processed to consolidate the sequences for different genome segments of each virus isolate and remove duplicated information. Then, GISAID and NCBI data were combined by matching isolate names and metadata, resulting in a consolidated dataset where each virus isolate was classified according to the datasets where it was represented (GISAID only, NCBI only, or both; S4 File). For each virus isolate, a whole genome equivalent (WGE) was calculated as the sum of the binary availability of sequence data for each gene segment (PB2, PB1, PA, HA, NP, NA, MP, NS) divided by eight. WGE ranged from 0.125 to 1; for example, WGE equal to 0.125 indicates that a sequence is available for only one gene segment of an isolate, whereas WGE equal to 1 indicates that sequences are available for all eight segments of an isolate.

In addition to WAHIS, EMPRES-i+, GISAID, and NCBI, we also compiled information on wildlife mortality and HPAI H5 cases from other public sources such as official government websites, scientific literature, and reports from government authorities, protected areas, and research groups. Additionally, internet searches were performed with Google, GoogleScholar and PubMed combining key terms (avian influenza, bird flu, influenza A, HPAI, H5, H5N1, mortality, outbreak, confirmed, suspected, bird, fauna, seal, wildlife) and the name of countries/territories in the Americas in four languages (English, Spanish, Portuguese, and French). Exceptionally, when no official or scientific sources could be located for a given mortality event or outbreak, information contained in news articles was also considered. The sources of information considered for each host-country association are explicitly provided in the “Source” column of the dataset (S1 File), and a reference list is provided (S2 File). Data were considered from public websites and databases as of 31 July 2025 and from scientific papers and preprints released online up to 31 December 2025.

### Data analysis

Through the consolidation and cross-referencing of the aforementioned data sources, a consolidated dataset was compiled on the publicly-reported wildlife found dead or estimated to have died in association with HPAI H5 in the Americas during the study period. The consolidated dataset was structured as a list of host taxa and country/territory combinations, with the following variables for each taxon-country association: (a) number of individuals publicly-reported to have been found dead or estimated to have died in association with HPAI H5 (based on the combined interpretation of all sources in the study), (b) number of individuals affected by HPAI H5 as represented in WAHIS, (c) H5 virus genome data availability (in whole genome equivalents; calculated separately for GISAID and NCBI as well as for the combination of both), and (d) whether or not EMPRES-i+ reported HPAI H5 for the taxon-country association.

The conservation status, English common name, and native/non-native status of each taxon relative to the country and to the Americas were compiled from the International Union for Conservation of Nature [76]. Junior taxonomic synonyms (i.e. other taxonomic names used to refer by some sources to refer to the same taxon) were also recorded. Furthermore, each taxon-country was classified as to the existence of at least one confirmed HPAI H5 detection (as of 31 July 2024) in relation to each country, in the Americas, and at the global level; this assessment was based on data compiled in this study as well as on global data from WAHIS and the Food and Agriculture Organization (FAO 2025). Only taxon-country combinations with at least one confirmed HPAI H5 detection at the national level were included in the analyses (i.e. “TRUE” in the column “Positive_Country” in S1 File).

Analyses and plots were done using *R* v4.4.1 [77] with packages *ggplot* v3.5.1 [78], *ggsankey* v0.0.99 [79], *RColorBrewer* v1.1-3 [80], and *scales* v1.4.0 [81]. The denominator of the representation ratio was calculated dividing the publicly-reported mortality (number of individuals) by the WAHIS-reported affected wildlife (number of individuals) or by the available H5 virus genomic data (whole genome equivalents). Zero-intercept linear models relating log_10_(y) to log_10_(x) were used to assess the correlation between variables.

## Supporting information

**S1 Fig. Correlation between the number of wildlife species with publicly-reported mortality associated with HPAI H5 and the number of wildlife species with HPAI H5 cases represented in WAHIS data (A, B), EMPRES-i+ data (C, D) or with available H5 virus genomic data (E, F) in the Americas**. The names of countries/territories (A, C, E) and host families (B, D, F) are abbreviated as in Tables 1 and 2 (host families with a single species affected are not labelled). Regression lines (and their 95% confidence interval in shaded areas) are drawn separately for wild birds (red) and wild mammals (blue) and regression parameters are shown above each plot. Study period: November 2021 – July 2024 (inclusive).

(PDF)

**S1 Table. Summary of the number of wildlife species associated with HPAI H5 in different datasets among countries/territories in the Americas**. Study period: November 2021 – July 2024 (inclusive).

(PDF)

**S2 Table. Summary of the number of wildlife species associated with HPAI H5 in different datasets among host families in the Americas**. Study period: November 2021 – July 2024 (inclusive).

(PDF)

**S1 File. Consolidated dataset on wildlife publicly-reported to have been found dead or estimated to have died in association with HPAI H5 in the Americas, November 2021 – July 2024**.

(CSV)

**S2 File. List of data sources**. (PDF)

**S3 File. Details of data processing and dataset compilation methods. (**PDF)

**S4 File. Merged GISAID/NCBI dataset of H5 virus genomic sequencing data from wildlife in the Americas, November 2021 – July 2024 (data available as of 31 July 2025)**.

(CSV)

### Acknowledgements

We would like to express our sincere appreciation to all professionals who have worked diligently and courageously, confronting the inherent risks of field and laboratory work, in their efforts to document and investigate wildlife mortalities associated with HPAI across the Americas. We also recognize the emotional and mental wellbeing challenges faced by those responding to large-scale wildlife mortality events, often under difficult, uncertain, resource-limited, and sensitive conditions. Their empathy, professionalism, and commitment to wildlife health are central to this work. This study was funded by University of California, Davis, with contributions from U.S. National Science Foundation Center for Pandemic Insights (NSF CPI, award number 241252) and CEVA Wildlife Research Fund.

## Author contributions

**Conceptualization:** Ralph E. T. Vanstreels, Marcela Uhart.

**Data curation and formal analysis:** Ralph E. T. Vanstreels.

**Investigation:** Ralph E. T. Vanstreels, Marcela Uhart.

**Writing – original draft:** Ralph E. T. Vanstreels.

**Writing – review & editing:** Ralph E. T. Vanstreels, Marcela Uhart.

## Notes

### Competing Interest Statement

The authors have declared no competing interest.

