## Supplementary material for "Data gaps of international databases on HPAI H5 in wildlife in the Americas: implications for surveillance, research, and conservation": S1 Table

**S1 Table. Summary of the number of wildlife species associated with HPAI H5 mortality in different datasets among countries/territories in the Americas.** Study period: November 2021 – July 2024 (inclusive).

| Country/territory | This study | WAHIS | EMPRES-i+ | GISAID | NCBI | GISAID+NCBI |
|---|---|---|---|---|---|---|
| **Wild birds** | | | | | | |
| Argentina (ARG) | 11 | 9 | 7 | 3 | 3 | 3 |
| Bolivia (BOL) | 1 | 1 | 1 | 0 | 0 | 0 |
| Brazil (BRA) | 23 | 23 | 13 | 9 | 9 | 9 |
| Canada (CAN) | 96 | 89 | 45 | 83 | 0 | 83 |
| Chile (CHL) | 58 | 54 | 19 | 28 | 24 | 28 |
| Colombia (COL) | 5 | 5 | 1 | 1 | 0 | 1 |
| Costa Rica (CRI) | 4 | 4 | 3 | 2 | 0 | 2 |
| Cuba (CUB) | 4 | 4 | 3 | 0 | 0 | 0 |
| Ecuador (ECU) | 7 | 5 | 4 | 2 | 0 | 2 |
| Falkland/Malvinas (FLK) | 7 | 6 | 4 | 1 | 0 | 1 |
| Greenland (GRL) | 3 | 3 | 1 | 2 | 0 | 2 |
| Guatemala (GTM) | 1 | 1 | 1 | 1 | 0 | 1 |
| Honduras (HND) | 1 | 1 | 1 | 1 | 0 | 1 |
| Mexico (MEX) | 18 | 15 | 5 | 0 | 1 | 1 |
| Panama (PAN) | 2 | 2 | 1 | 1 | 0 | 1 |
| Peru (PER) | 39 | 30 | 18 | 10 | 8 | 11 |
| United States (USA) | 177 | 161 | 19 | 127 | 115 | 128 |
| Uruguay (URY) | 7 | 5 | 3 | 2 | 2 | 2 |
| Venezuela (VEN) | 2 | 1 | 0 | 2 | 1 | 2 |
| **Total** | 287 | 261 | 105 | 174 | 142 | 175 |
| **Wild mammals** | | | | | | |
| Argentina (ARG) | 3 | 3 | 3 | 3 | 3 | 3 |
| Brazil (BRA) | 2 | 2 | 2 | 1 | 1 | 1 |
| Canada (CAN) | 11 | 7 | 5 | 8 | 0 | 8 |
| Chile (CHL) | 7 | 5 | 6 | 5 | 6 | 6 |
| Peru (PER) | 3 | 2 | 1 | 3 | 3 | 3 |
| United States (USA) | 26 | 21 | 18 | 14 | 10 | 14 |
| Uruguay (URY) | 3 | 3 | 3 | 3 | 3 | 3 |
| **Total** | 39 | 31 | 27 | 26 | 20 | 26 |
