## Supplementary material for "Data gaps of international databases on HPAI H5 in wildlife in the Americas: implications for surveillance, research, and conservation": S2 Table

**S2 Table. Summary of the number of wildlife species associated with HPAI H5 mortality in different datasets among host families in the Americas.** Study period: November 2021 – July 2024 (inclusive).

| Country/territory | This study | WAHIS | EMPRES-i+ | GISAID | NCBI | GISAID+NCBI |
|---|---|---|---|---|---|---|
| **Wild birds** | | | | | | |
| Accipitridae (Acc) | 19 | 19 | 12 | 14 | 11 | 14 |
| Alcidae (Alc) | 5 | 5 | 3 | 4 | 1 | 4 |
| Anatidae (Ana) | 67 | 60 | 21 | 44 | 35 | 44 |
| Anhimidae (Anh) | 1 | 0 | 0 | 0 | 0 | 0 |
| Apodidae (Apo) | 1 | 0 | 0 | 1 | 1 | 1 |
| Ardeidae (Ard) | 8 | 7 | 2 | 5 | 5 | 5 |
| Bombycillidae (Bom) | 1 | 0 | 0 | 0 | 0 | 0 |
| Cardinalidae (Car) | 1 | 0 | 0 | 0 | 0 | 0 |
| Casuariidae (Casu) | 1 | 1 | 0 | 1 | 1 | 1 |
| Cathartidae (Cat) | 5 | 5 | 4 | 4 | 3 | 4 |
| Charadriidae (Cha) | 4 | 4 | 1 | 2 | 2 | 2 |
| Ciconiidae (Cic) | 1 | 1 | 0 | 1 | 1 | 1 |
| Columbidae (Col) | 4 | 3 | 2 | 2 | 1 | 2 |
| Corvidae (Cor) | 6 | 6 | 2 | 5 | 4 | 5 |
| Diomedeidae (Dio) | 2 | 2 | 2 | 0 | 0 | 0 |
| Estrildidae (Est) | 1 | 1 | 0 | 0 | 0 | 0 |
| Falconidae (Fal) | 8 | 7 | 2 | 6 | 5 | 6 |
| Fregatidae (Fre) | 2 | 2 | 2 | 1 | 1 | 1 |
| Fringillidae (Fri) | 2 | 0 | 0 | 0 | 0 | 0 |
| Gaviidae (Gav) | 2 | 2 | 0 | 1 | 1 | 1 |
| Gruidae (Gru) | 2 | 2 | 0 | 1 | 1 | 1 |
| Haematopodidae (Hae) | 2 | 2 | 1 | 1 | 1 | 1 |
| Hirundinidae (Hir) | 3 | 3 | 2 | 0 | 0 | 0 |
| Hydrobatidae (Hyd) | 1 | 0 | 0 | 0 | 0 | 0 |
| Icteridae (Ict) | 4 | 3 | 1 | 2 | 2 | 2 |
| Laridae (Lar) | 31 | 31 | 16 | 26 | 26 | 27 |
| Oceanitidae (Oce) | 1 | 1 | 0 | 0 | 0 | 0 |
| Odontophoridae (Odo) | 1 | 1 | 0 | 0 | 0 | 0 |
| Pandionidae (Pan) | 1 | 1 | 0 | 1 | 1 | 1 |
| Parulidae (Par) | 0 | 0 | 0 | 0 | 0 | 0 |
| Passerellidae (Pas) | 3 | 3 | 0 | 0 | 0 | 0 |
| Pelecanidae (Pel) | 4 | 4 | 3 | 3 | 3 | 3 |
| Phaethontidae (Phae) | 1 | 0 | 0 | 0 | 0 | 0 |
| Phalacrocoracidae (Phal) | 7 | 6 | 4 | 3 | 3 | 3 |
| Phasianidae (Phas) | 4 | 3 | 0 | 3 | 1 | 3 |
| Phoenicopteridae (Phoe) | 3 | 3 | 0 | 0 | 0 | 0 |
| Platylophidae (Pla) | 1 | 0 | 0 | 1 | 0 | 1 |
| Podicipedidae (Pod) | 6 | 6 | 2 | 5 | 5 | 5 |
| Procellariidae (Proce) | 10 | 10 | 3 | 5 | 2 | 5 |
| Psittacidae (Psici) | 11 | 11 | 0 | 1 | 1 | 1 |
| Psittaculidae (Psicu) | 0 | 0 | 0 | 0 | 0 | 0 |
| Rallidae (Ral) | 2 | 2 | 1 | 1 | 1 | 1 |
| Ramphastidae (Ram) | 2 | 2 | 0 | 0 | 0 | 0 |
| Rheidae (Rhe) | 1 | 1 | 0 | 0 | 0 | 0 |
| Scolopacidae (Sco) | 10 | 9 | 3 | 9 | 8 | 9 |
| Spheniscidae (Sph) | 4 | 2 | 2 | 1 | 1 | 1 |
| Stercorariidae (Ste) | 4 | 4 | 2 | 2 | 1 | 2 |
| Strigidae (Stri) | 12 | 12 | 4 | 10 | 8 | 10 |
| Struthionidae (Stru) | 1 | 1 | 1 | 0 | 0 | 0 |

| Country/territory | This study | WAHIS | EMPRES-i+ | GISAID | NCBI | GISAID+NCBI |
|---|---|---|---|---|---|---|
| Sturnidae (Stu) | 1 | 1 | 1 | 0 | 0 | 0 |
| Sulidae (Sul) | 6 | 5 | 5 | 4 | 3 | 4 |
| Threskiornithidae (Thr) | 5 | 5 | 1 | 2 | 1 | 2 |
| Turdidae (Tur) | 1 | 1 | 0 | 1 | 1 | 1 |
| Tyrannidae (Tyr) | 1 | 1 | 0 | 1 | 0 | 1 |
| **Total** | 287 | 261 | 105 | 174 | 142 | 175 |
| **Wild mammals** | | | | | | |
| Canidae (Can) | 3 | 2 | 2 | 2 | 1 | 2 |
| Castoridae (Cast) | 1 | 0 | 0 | 0 | 0 | 0 |
| Cricetidae (Cri) | 3 | 1 | 2 | 0 | 0 | 0 |
| Delphinidae (Del) | 4 | 2 | 3 | 4 | 3 | 4 |
| Didelphidae (Did) | 1 | 1 | 1 | 1 | 0 | 1 |
| Felidae (Fel) | 6 | 6 | 3 | 3 | 2 | 3 |
| Leporidae (Lep) | 1 | 1 | 1 | 0 | 0 | 0 |
| Mephitidae (Mep) | 1 | 1 | 1 | 1 | 1 | 1 |
| Mustelidae (Mus) | 6 | 6 | 4 | 4 | 3 | 4 |
| Otariidae (Ota) | 2 | 2 | 2 | 2 | 2 | 2 |
| Phocidae (Phoci) | 3 | 3 | 2 | 3 | 2 | 3 |
| Phocoenidae (Phoco) | 2 | 0 | 1 | 1 | 1 | 1 |
| Procyonidae (Procy) | 2 | 2 | 2 | 2 | 2 | 2 |
| Sciuridae (Sci) | 1 | 1 | 0 | 0 | 0 | 0 |
| Ursidae (Urs) | 3 | 3 | 3 | 3 | 3 | 3 |
| **Total** | 39 | 31 | 27 | 26 | 20 | 26 |
