## Supplementary figures and images for "Data gaps of international databases on HPAI H5 in wildlife in the Americas: implications for surveillance, research, and conservation"

### S1 Fig

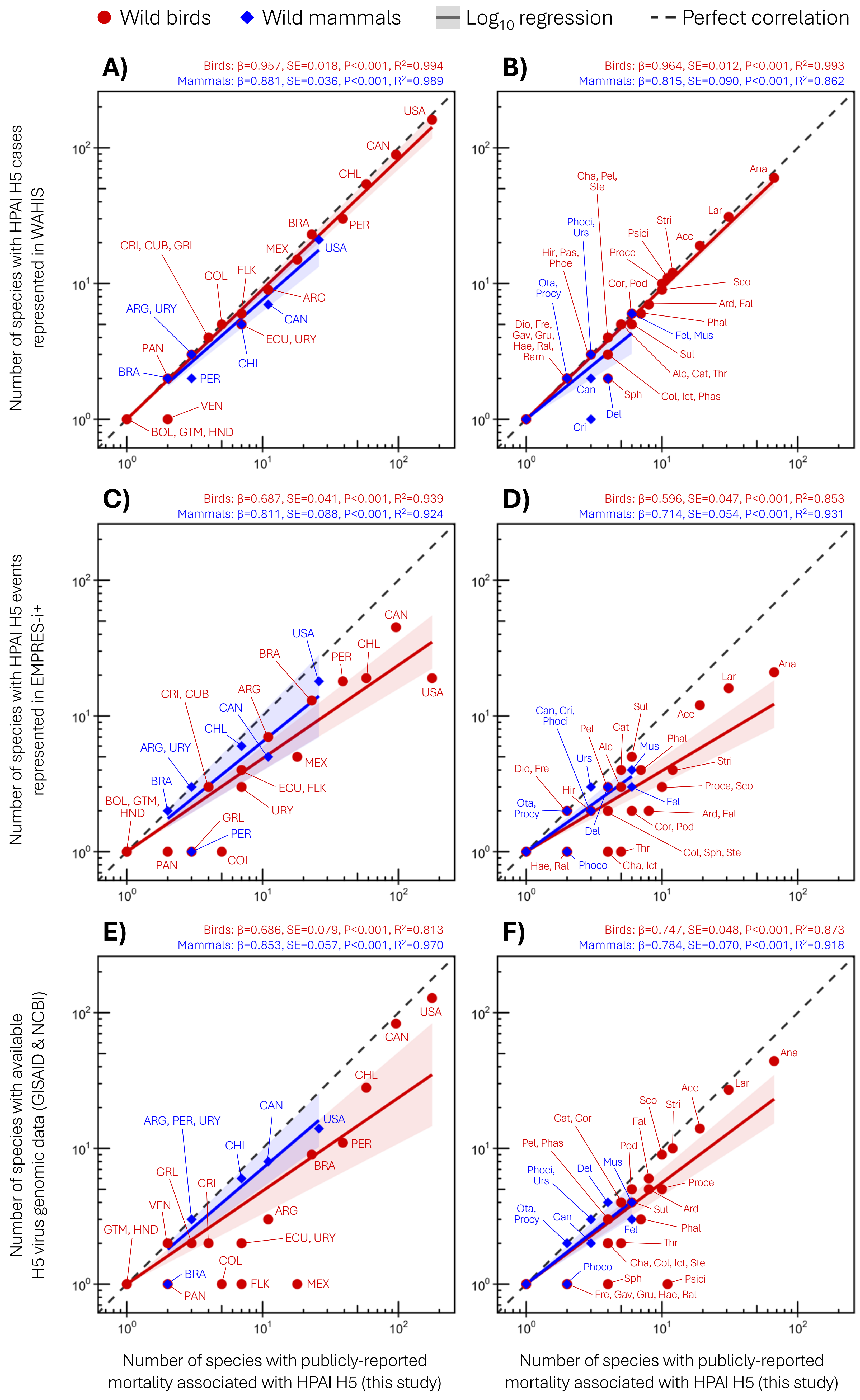
